# Biochemical and subcellular characterization of a squid hnRNPA/B-like protein in osmotic stress activated cells reflects molecular properties conserved in this protein family

**DOI:** 10.1101/2021.07.02.450876

**Authors:** Gabriel S. Lopes, Diego T. P. Lico

## Abstract

In previous works, we characterized a novel, strongly basic, squid hnRNPA/B-like Protein 2 in presynaptic terminals of squid neurons. Here, we show that squid hnRNPA/B-like Protein 2 are exclusively nuclear localization and relocated to cytoplasmic granules containing hnRNPA1 and Poly-A binding protein-1 (PABP-1) when the cells are treated with sorbitol. Also, we show an interaction of hnRNPA/B like Protein 2 with squid RNA, its interfered with dynamic of formation of hnRNPA/B like Protein 2 dimers, whereas possibly involved disulfide bounds and postranslations modification in a distinct stage of dimers formation. An understanding of the molecular and biochemical mechanisms involved in the stability of the dimeric form, and the regulation of the transition between monomeric and dimeric forms may bring insights into evolution of several neurodegenerative diseases.

**Highlights:**

- We identified monomeric (p37) and dimeric (p65) forms of squid hnRNPA/B-like Protein 2 in squid optic lobes
- Our data indicate a conserved structure and cellular properties of squid hnRNPA/B-like Protein 2 and human hnRNPA1 protein colocalizing with PABP into stress granules (SGs)
- The stability of hnRNPA/B-like dimers involved the squid RNAs and disulfide bonds to promote higher SDS-stable dimers formation
- An understanding of the transition between monomeric and dimeric forms of squid hnRNPA/B-like Protein 2 may give clues to misfolding processes in neuropathologies

## Introduction

Two hnRNPA/B-like proteins were identified associated with tissue purified squid p65. We hypothesized that squid p65 is an SDS-stable dimer composed of two ~37-kDa subunits: hnRNPA/B-like Protein 1 (36.3 kDa) and Protein 2 (37.6 kDa), Lico et al., 2015.

Members of hnRNPA/B proteins display multiple functions and are structurally conserved family of RNA-binding proteins, that form core elements in ribonucleoprotein complexes involved in packaging of nascent mRNAs into granules, can shuttle out of the nucleus relocated to cytoplasmic (for reviews, see Siomi H et al., 1995; Michael WM et al., 1995. Dreyfuss G et al., 2002; He & Smith 2009 and McDonald et al., 2011). Notably, hnRNP A1 and hnRNP A2B1 are components of RNA transport granules in neurons (Heraviab Y. B. et al., 2020, Lopes et al 2020, Purice MD & Taylor JP 2018, Douglas J N et al., 20016, Lico et al., 2015 and Elvira et al., 2006).

Here, we show cellular properties conserved between squid hnRNPA/B-like Protein 2 and human hnRNPA1 protein in neuroblastoma cells (SH-SY5Y), where squid hnRNPA/B-like Protein 2 shuttle out of the nucleus relocated to cytoplasmic into granule containing hnRNPA1 and Poly-A binding protein-1 (PABP-1). Following the dynamic from stress granules (SGs) undergo fusion, fission and flowing in the cytosol, as Guil S. et al., 2006

Squid hnRNPA/B-like Protein 2 and human hnRNP A1 protein have primary structure conserved with two RNA binding domains or RRM domains in the N-terminal half of molecules (Lopes et al., 2019), in which showed a selectively RNA interaction by squid hnRNPA/B-like protein 2 at presynaptic terminal of squid neuron (Lopes et al., 2020 and Maris C et al., 2005). These domains are sites of interaction with RNA molecules formed by disulfide bonds between RNA binding domains in a poly(A)-binding protein can regulate the ability of this protein to bind mRNA (Fong C. L et al., 2000).

Many are studies aimed at determining the assembly mechanisms of mRNP granules have focused on proteins. However, in a recent study, suggested an ability of RNA self-assembly that could contribute to granules formation in cytoplasm; e.g. secondary structure of RNA could act regulating binds of specific RNA and determines their interaction with other RNAs molecules or with RNA binding proteins into granules cytoplasmic of cells (see Khong A et al., 2017; Maharana et al., 2018, Langdon et al., 2018, Van Treeck B & Roy Parker, 2018 and Sterneckert J et al., 2018). Thus, the propensity to aggregate reversibly is an important property for components of granules which can be regulatable by protein–protein, protein–RNA and RNA-RNA interaction. These control of key components in the granules formation can induce a shift from repressive to active translational function of mRNAs. Uncontrolled disrupt their normal function can be deregulating RNA processing and translation in the cells. However, its not clear exactly, what ungoverned aggregation could lead the formation of inclusion bodies in the several neurodegenerative diseases (Zhang K et al., 2018 and Yalda B.H.et al., 2020, respectively)

A variety of neurodegenerative diseases are associated to mRNPs mutation or post-translational modifications like phosphorylation, ubiquitination, acetylation, cysteine oxidation, and sumoylation (Buratti et al., 2018). In particular, human hnRNP A1-and A2/B1 harbor a steric-zipper-motif in the C-terminal domain that is affected by identified mutations and which accelerates self-seeding fibril formation and cross-polymerization with wild-type protein (Kim HJ et al., 2013; Kwiatkowski Jr. et al., 2009; Liu, Y.C et al., 2013; Liu, Q et al., 2016, Mackenzie et al., 2017; Protter, D.S.W et al., 2018; Meyer-Luehmann et al., 2020; Aguzzi and Rajendran, 2009), in which has leads the formation of inclusion bodies in the several neurodegenerative diseases. Finally, there are, many unanswered questions surrounding the role hnRNPs play in neurodegenerative diseases.

### Experimental Procedures

#### Brain tissue preparation

Optic lobes were dissected from *Dorytheuthis pealeii* obtained from the Marine Resources Center of the Marine Biological Laboratory in Woods Hole, MA and from *Dorytheuthis plei* obtained from the Centro de Biologia Marinha-CEBIMar, University of São Paulo, São Sebastião, Brazil. Brains tissues were quickly frozen in liquid nitrogen and stored at −70°C until used. Homogenated tissues (~100 mg each) were prepared in a IKA^®^ RW 20 digital homogenizer in phosphate buffer (50 mM sodium phosphate, pH 7,5, 100 mM NaCl, 25 mM DTT), 1 ml/sample, containing a cocktail of protease inhibitors (1 mM benzamidine, 1 mM AEBSF and 1 μg/ml aprotinin) and then centrifuged at 50.000 x g for 30 min. at 4°C. The supernatant 1 (S1) was maintained on ice until use.

#### Preparation of total RNA from squid optic lobes

Total RNA from squid optic lobes was prepared as described in Chomczynski and Sacchi (1987) with slight modifications. Briefly, one optic lobe (~100 mg) was homogenized in 1.2 ml of TRIzol reagent in a glass-teflon homogenizer at room temperature (RT). The homogenate was centrifuged at 10,200 × g for 10 min. at 2-8°C (Eppendorf, refrigerated microfuge). One ml of supernatant (S1) was removed avoiding the lipid overlayer, transferred to a clean microfuge tube and incubated at R.T. for 5 min. Then, 200 μl of chloroform were added, mixed vigorously by inversion, vortexed for 15 seconds and incubated at R.T. for 2 min. The tube was centrifuged at 10,200 x g for 20 min at 4°C. The upper aqueous phase (~600 ul – containing RNA) was carefully transferred to a transparent microfuge tube, and RNA was then precipitated by addition of an equal volume of chilled isopropanol. The tube was incubated in the freezer for at least 2 h to overnight, and then centrifuged at 10,200 x g for 30 min at 4°C. The supernatant was carefully removed. The RNA was visualized as a gel pellet in the bottom of the microfuge tube. The RNA gel was washed twice with 1 ml of ice cold 70 % ethanol by light vortexing and centrifuging at 10,200 x g for 10 min at 4°C. RNA was stored in 70% ethanol for a week at 4°C or for a few months at −20°C. RNA gel was air dried until ethanol completely evaporate (10-20 min). RNA was dissolved in 100 μl of DEPC water, and then treated with 10 μl DNase I (Fermentas, 1U/ul) for 15 min at 37°C. The reaction was stopped by extraction in phenol/chloroform or TRIzol. RNA was dissolved in 100 μl of DEPC treated water.

#### Biochemical assay

*Urea-PAGE*: minigels were prepared according to McLellan, 1982, with slight modifications (Lico et al., 2015). *Purification by cationic and reverse phase chromatography*: Purification of p65 and p37 or *Affinity purified squid hnRNPA/B-like Protein 2* in fusion with a 6xHis tag was expressed in bacteria and purified as described was done as previously described (Lico et al., 2010 and 2015, respectively). *Gel filtration chromatography* sample was fractionated by fast protein liquid chromatography (FPLC) (äKTA purifier, GE Health Science) was performed using Superdex 75 3.2/300 column in a phosphate buffer with 4 M urea in the presence of 125 mM of DTT. For calibration of gel filtration column, rabbit IgG (Mr 120,000, affinity purified), Bovine serum albumin (Mr 66,000, Sigma) and hydrogen peroxide (H2O2, Merck) were used.

#### DNA constructions

Full-length squid hnRNPA/B-like Protein 2 was amplified by PCR using full-length squid cDNA and corresponding primers, cloned into pGEM-T Easy vector (Promega Corporation, Wisconsin, USA), and subcloned into pEGFP-C1 and pmCherry-C1 vectors (Clontech) for expression in mammalian cells. Squid hnRNPA/B-like Protein 2 transcript was amplified using the folowing forward and reverse primers, respectively: 5’–AT<u>GAATTC</u>TATGCCCGAAAGGTAC–3’ and 5’AT<u>GGGCCC</u>**TTA**CCGTCTGTAACCGCC-3’. *EcoR* and *ApaI* restriction sites are underlined. The complementary stop codon for the C-terminus is shown in bold. DNA was sequenced using the Big Dye Terminator Cycle Sequencing Ready Reaction (Applied Biosystems, Foster City, CA, USA) on ABI 3100 sequence analyzer. Affinity purified squid hnRNPA/B-like Protein 2 fused to a 6xHis tag was expressed in bacteria and purified as previously described (Lico et al., 2015). Using the same restriction enzimes, for expression in drosophila S2 cell line, DNA fragment encoding squid Protein 2 was cloned in pAc 5.1-B vector.

#### Cell culture and transfection

Human SH-SY5Y neuroblastoma cell line was a Kindly gift of Dr. Adriano Sebollela (Biochemistry Department of University of São Paulo). SH-SY5Y neuroblastoma cell line was grown in Dulbecco’s modified Eagle’s medium (DMEM) high glucose (Sigma-Aldrich St. Louis, MO, USA) supplemented with penicillin (100 units/ml), streptomycin (100 μg/ml), and 10% (vol/vol) heat-inactivated fetal bovine serum (FBS) (GIBCO, Gaithersburg, MD, USA). Cells were maintained at 37°C in a saturated humidity atmosphere containing 95% air and 5% CO2. For SH-SY5Y cells transfection, 9 × 10 cells/cm^2^ were plated on 13 mm^2^ glass slides, in a 24 well plate. 24 hours later, cells were transfected for 3 h using 0.5 μg per well of pEGFP-C1 or pmCherry-C1 vectors (containing or not the DNA constructs), and 1 μl lipofectamine 2000 following manufacture instructions (Invitrogen, Life Technologies, Grand Island, NY, USA). After incubation, transfection medium was replaced by DMEM supplemented with penicillin (100 units/ml), streptomycin (100 mg/ml), and 10% FBS. Non-adherent, embryonic *Drosophila* cells (S2) were grown in Schneider media (Invitrogen, Carlsbad, CA, USA) containing 10% heat-inactivated fetal bovine serum (FBS), antibiotic/antimycotic solution (Invitrogen), and maintained at 26°C in a saturated humidity atmosphere without CO2. S2 cells were plated 24 hours before transfection in a 12 well plate (1 x 10^6^ cells per well). In the day after, transfection was carried out using *Effectene Transfection ReagentKit* (Qiagen, Maryland, USA), following manufacture instruction, and 0.5 μg of *pAc5.1-B* vector containing the open reading frame (ORF) of squid hnRNPA/B-like Protein 2 was used. **Osmotic stress.** D-Sorbitol (Sigma) was diluted in standard growth medium to yield a 0.4 M concentration. After 1 h of incubation in this medium, cells were fixed and processed for immunostaining.

#### Immunocytochemistry

After 48 h post-transfection, cells were fixed and permeabilized in phosphate-buffered saline (PBS) containing 2% paraformaldehyde and 0.3% Triton X-100 for 20 min, then blocked with 2% Bovine Serum Albumin (BSA) in PBS for 1 h, washed 3x times with PBS, and incubated with 10 μg/ml monoclonal anti-PABP and/or 6 μg/ml anti-hnRNPA1 diluted in PBS containing 1% BSA, for 2 h at RT in a humid chamber. Primary antibodies were visualized with secondary antibody conjugated with Alexa Fluor 488, 594 or 647 (Molecular Probes, Invitrogen).

Secondary antibodies waere incubated for 1 h diluted in PBS containing 1% BSA. Nuclei were detected using DAPI. Stained cells were examined using a confocal microscope TCS-SP5 (*Leica Microsystems*, Germany), or an inverted microscope *LSM 780 AxioObserver* (*Carl Zeiss*, Germany).

#### Fluorescence Recovery After Photobleach (FRAP)

About 1 x 10^5^ cells were plated on 14 mm^2^ coverslip and left in a humid chamber at 37°C / 5% CO_2_ to adhere. About 24 h later, cells were transfected with the pEGFP-C1 vector containing the DNA insert encoding the recombinant hnRNPA/B-like Protein 2 (rP2). Approximately 20 hours after transfection cells were treated, or not (control), with 400 mM D-Sorbitol diluted in DMEM medium containing 10% SFB and 1% Penicillin / Streptomycin solution and left in the humid chamber at 37°C / 5% CO_2_. After one hour, living cells were analyzed by FRAP in an inverted microscope LSM 780 AxioObserver (Carl Zeiss), where the post bleach recovery rate of cytoplasmic stress granules were determined. The microscope software (Zen Black) was set to scan the region of interest 10 times (one second each) before bleaching. Variations in fluorescence intensity of the bleached region were followed for up to 180 seconds, and values recorded every second. To calculate the recovery percentage, which represents the portion of moving elements in the region of interest (ROI), we used the following formula: percentage = 100x [(IFR-IFB) / (IFPB-IFB)], where IFR = recovered fluorescence intensity; IFB = fluorescence intensity achieved in bleach; IFPB = pre-bleach fluorescence intensity. To calculate recovery kinetics, the following formula was used: Ʈ1 / 2 = tR½-t0, where Ʈ1 / 2 = the recovery half-life; t0 = the time when the bleach was maximum; tR½ = the time it takes for recovery to reach half of its final value.

#### Western Blotting

SDS-PAGE was performed on 10% minigels (BioRad, Hercules, CA). After electrophoresis, proteins were transferred to nitrocellulose membranes. After incubation with anti-sqRNP2 antibody, a secondary anti-rabbit HRP antibody, membrane was developed using ECL^™^ Western Blot Detection Reagent (GE Healthcare, RNP2209) following manufacturer instructions.

#### Antibodies

Monoclonal anti-hnRNPA1 (Sigma, clone 9H10, n° R4528) and anti-PolyA-Binding Protein (anti-PABP) (Sigma, clone 10E10, n° P6246) were commercially acquired. Anti-squid RNP2 antibody (α-sqRNP2) was raised in rabbits against a synthetic peptide CLFIGGLSYDTNEDTIKK (BioSynthesis, Lewisville, Texas), as described (Lico et al. 2015).

## Results

We proposed p65 to be an SDS-stable dimer composed of ~37-kDa hnRNPA/B-like subunits. In this report, we have investigated new details of molecular properties involved in the stability of dimeric form, and their regulation in the transition between monomeric and dimeric forms of hnRNPA/B-like protein 2. In which, its could be involves disulfide bounds and their ability of bind RNA, to transform into a urea-stable tetramer and SDS-stable dimer as is characteristic of tissue purified p65.

Thus, the mobility of bacterially expressed hnRNPA/B-like protein 2 (rP2), corresponds to cellular p37, was investigated by gel filtration chromatography. We show the higher oligomers of rP2 (with ≥80-kDa) in urea solution, which included the formation of dimers that are not stable as in SDS-PAGE. Whereas when treated with high concentrations of DTT (125 mM DTT in phosphate buffer), there was a disrupted their higher oligomers of rP2, in two distinct fractions (≥ 80-kDa and ~40-kDa) can be visualized by Urea–PAGE (Fig. 1a and 1b). However, high concentrations of DTT does not affected the mobility of endogenous p65 (Fig. S3). Remarkably, the typical SDS-stable dimers were formed with the amount increase of squid optic lobe extract (Fig. 1c), and when there was raising to 125 mM of DTT included in the samples (Fig. 2d) by *in vitro* studies. Note, it was not observed with 25 mM of DTT in phosphate buffer visualized by immunoblotting.

**Figure 1.**
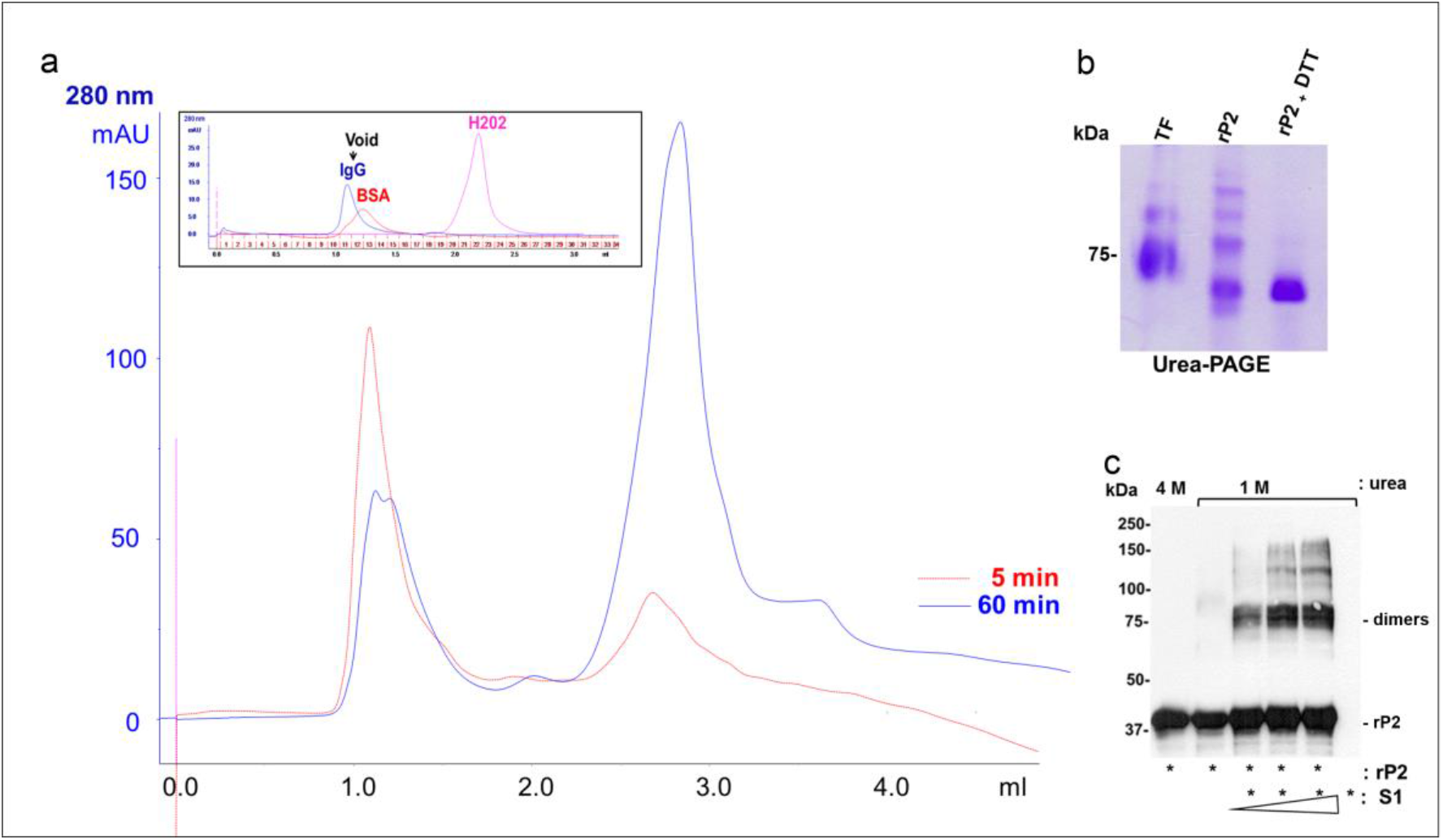
Biochemical properties of squid hnRNPA/B-like Protein 2 as a potential SDS-stable dimer. a) The bacterially expressed hnRNPA/B-like protein 2 (rP2) sample was fractionated by gel filtration chromatography (Superdex 75 3.2/300 column) on ÄKTA purifier systems in a phosphate buffer, 7.4 pH, containing 4 M urea in the presence of 125 mM DTT. After 5 minutes of incubation with DTT the major part of rP2 (1.4 mg/ml) fall into the void in 1.1 ml volume (red lane, peak: BSA/IgG, upper insert) and small peak in 2.7 ml volume (peak: H2O2, upper insert). On the other hand, after 60 minutes, the major part of rP2 fall into the void in 2.7 ml volume (blue lane, peak: H2O2). For calibration of gel filtration column, we used rabbit IgG (Mr 120,000) and BSA (Mr 66,000, Sigma) and hydrogen peroxide (H2O2, Merck) b) The disturb of the higher oligomers of recombinant hnRNPA/B-like protein 2 (rP2) into monomeric form. Urea–PAGE of the peak rP2 fraction from the gel filtration chromatography column stained with Colloidal Coomassie Blue (CCB). Lane rP2, recombinant hnRNPA/B-like Protein 2 (10 μg) and lane rP2+DTT, with addition 125 mM of DTT. Lane TF, Transferrin (10 μg), applied a basic-protein indicator (~75 kDa) c) The endogenous factor(s) induced posttranslational modifications in recombinant hnRNPA/B-like protein 2 (rP2) that promote the formation of the SDS-stable dimer *in vitro*. Western blot performed on 10% gel probed with α-sqRNP2 antibody. Lane 1, 62 μg/ml of rP2 in the presence of 4 M urea; lane 2, rP2 in the presence of 1 M urea; lane 3-5, rP2 in the presence of 110, 220, 440 μg/ml squid optic lobe extracts in 1 M of urea and lane 6, 110 μg/ml squid optic lobe extract alone in phosphate buffer. All samples were kept at room temperature for 16h

**Figure 2.**
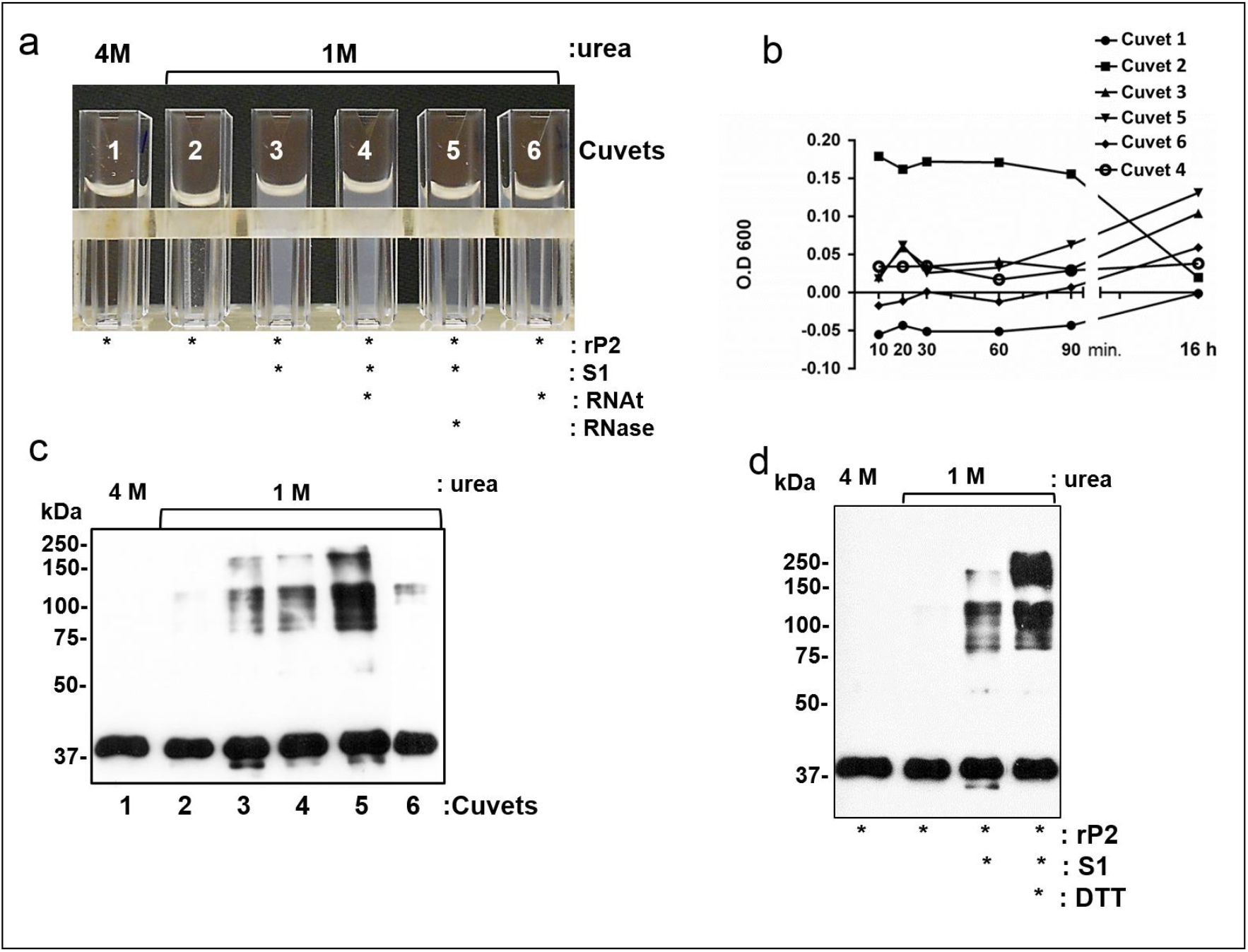
Formation of SDS-stable oligomers of hnRNPA/B-like Protein 2 can be affected by RNA and DTT *in vitro* studies. a) The visual analyses of precipitation of recombinant hnRNPA/B-like protein 2 (rP2) in 1 M urea over a 16 hours incubation period at room temperature (r.t). All cuvets contain 200 μg/ml of rP2 in 50 mM Phosphate Buffer, pH 7.5, 100 mM NaCl, 25 mM DTT, 30 mM Imidazole, 0,5 mM ATP and 1 mM protease inhibitors (aprotinin; Benzamidine and pefabloc). Cuvet 1 with 4 M urea; cuvet 2 with 1 M urea; cuvet 3 with 1 M urea and 220 μg/ml of supernatant 1 (S1) from squid optic lobe protein extract; cuvet 4 with 1 M urea and S1 plus 25 μg/ml of squid total RNADNA free; cuvet 5 with 1 M urea and S1 pre-treated with 10 μg/ml of RNase A by 30 minutes; and cuvet 6 with 1 M urea containing 25 μg/ml of squid total RNADNA free b) Chromatogram of the precipitation rP2 in low urea solution in all condition described above, which was accompanied by absorption at 600 nm for 16 hours at room temperature (r.t) c) The squid RNA molecules affected the formation of higher oligomers of rP2 *in vitro* studies. Western blot of samples mentioned above probed with polyclonal antibody α-sqRNP2 d) The reduction of disulfide bonds between RRM domains of rP2 and squid RNAs affected the formation of higher oligomers of rP2 *in vitro* studies. Western blot probed with α-sqRNP2 of samples carried out directly in plastic cuvets in 50 mM Phosphate Buffer, pH 7.5 as indicated in Fig. S2. All samples contain 200 μg/ml of rP2 in 50 mM Phosphate Buffer, pH 7.5, 100 mM NaCl, 25 mM DTT, 30 mM Imidazole, 0,5 mM ATP and 1 mM protease inhibitors (aprotinin; Benzamidine and pefabloc). Asterix indicated the addition of 220 μg/ml of S1 from squid optical lobe in and plus 125 mM of DTT in Phosphate Buffer

### Effects of RNA on the SDS-stable dimers

The precipitation of recombinant hnRNPA/B-like protein 2 (rP2) when diluted from 4 M to 1 M urea concentration can be verified by light scattering at 600 nm over a 16 h period at room temperature (Lico et al., 2015).

Here, to assess the effect of RNA in the precipitation of rP2. We isolated total RNA (RNAt) of squid optic lobe extract fraction. One total RNA fraction was pre-incubated at 95°C for 5 minutes to denaturation the RNA structures, other fraction was maintained at room temperature (r.t). In addition, the optic lobe extract fraction was treated with RNase-A to be incubated with rP2 *in vitro* experiments (Fig S1a and b).

When diluted from 4 M to 1 M urea concentration, the rP2 visibly precipitated from solution, which was accompanied by a rapid rise in absorption at 600 nm within minutes due to light scattering (Fig. 2a and b, cuvet 2). However, with addition of squid optic lobe extract (S1) or S1 plus total RNA, its inhibited precipitation preventing formation of flocculates and keep the rP2 soluble in low urea solution (Fig. 2a and b, cuvets 3 and 4, respectively). On the other hand, when the samples were examined by western blot after 16 h of incubation, the rP2 fraction treated with RNase-A formed higher SDS-stable dimers immunoreactive bands in the range of 80-250 kDa (Fig. 2c, cuvet 5). It was not verified in fraction containing only total RNA (RNAt /1 M) (Fig. 2c, cuvet 6), suggesting that squid RNA can capture rP2-binding and keep rP2 soluble in low urea solution, and endogenous factor(s) in the squid optic lobe extract produce post translational alterations in rP2 that induce its dimerization. All preparation of total RNA was verified the preservation of the RNA ribosomal by electrophoreses gels (data not show).

These data together suggests that squid RNAs and disulfide bonds could act as curb on availably of rP2 monomers to promote formation of higher SDS-stable dimers, whereas endogenous factor(s) in the squid optic lobe extract produce post translational alterations in rP2 that induce its dimerization, giving rise to a stable form equivalent to p65.

To searching information of squid hnRNPA/B-like Protein 2 that could regulated its specific oligomers; the endogenous p65 and p37 polypeptides were isolated of squid-tissue and further labeled with a phospho-specific dye (Pro-Q^®^ dye, Invitrogen). The tissue-purified p65 was labeled while p37 was not, showing phosphorylation distinct forms (Fig. 3c). Note, that rP2 corresponds to cellular p37 was not labeled. The sequence alignment of squid hnRNPA/B-like protein 2 and human hnRNP A1 protein primary structure; we observed potential phospho aminoacids in the hnRNP A1 protein has 42 Serine, 12 Threonine, and 19 Tyrosine residues of aminoacids, whereas squid hnRNPA/B-like protein 2 showed 14; 13 and 19, respectively. Our attention was caught by the differences of Serine numbers, and Cysteine (2/4); aromatic (20/26) and charged (14/18) residues of aminoacids between A1 and A/B-like proteins, respectively. In addition, human hnRNP A1 protein and squid hnRNPA/B-like Protein 2 showed structures conserved in critics domains which uncontrolled it accelerates self-seeding fibril formation and showed by incorporation of Thioflavin T fluoresces (ThT) into amyloid fibrils type (Fig. 3a-b and Kim HJ et al., 2013).

**Figure 3.**
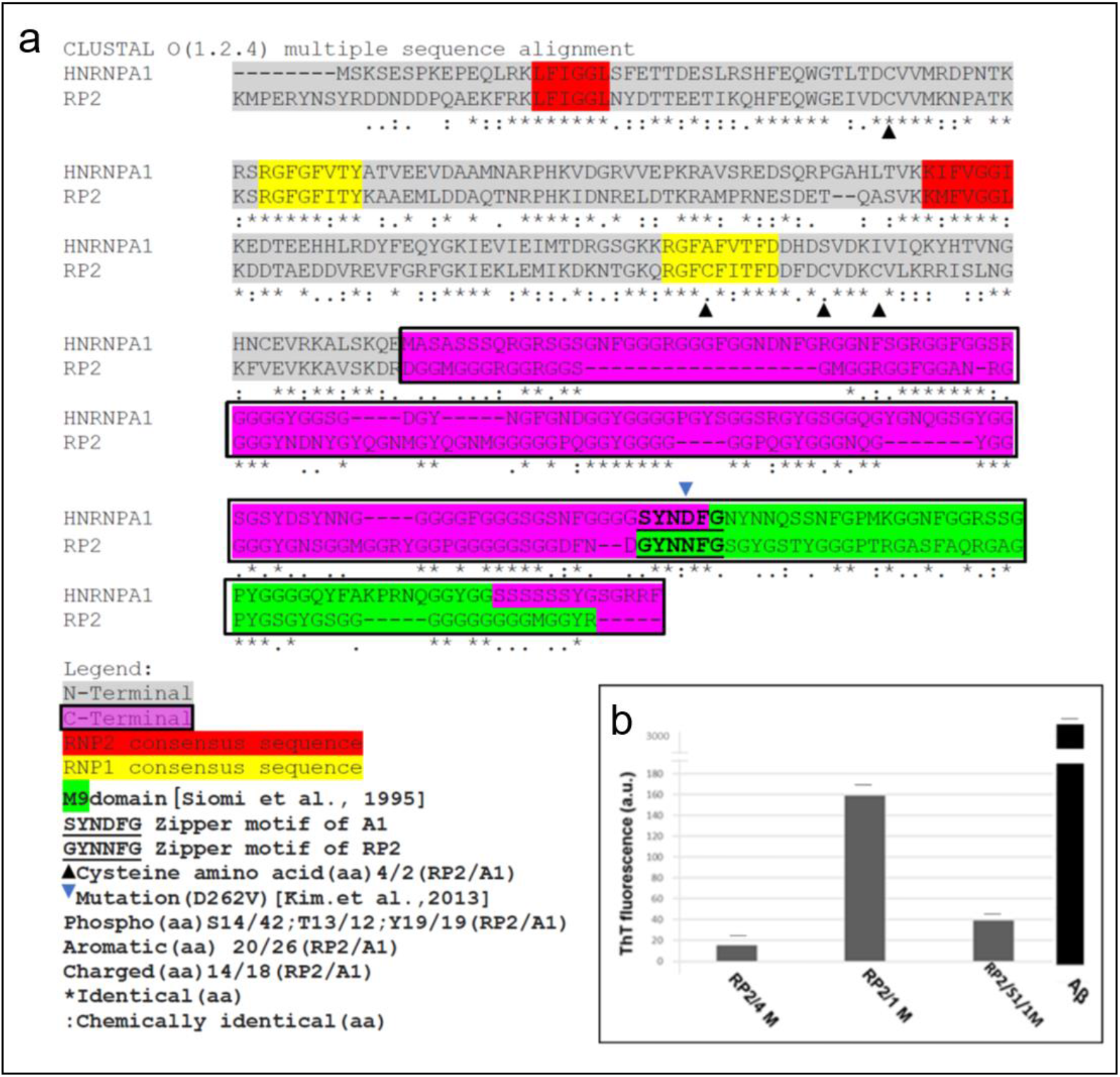
Sequence alignment of squid hnRNPA/B-like Proteins 2 primary structure having a potential *steric zipper motif* in the C-terminal, and a distinct phosphorylation between dimeric (p65) and monomeric (p37) forms tissue-purified. a) Sequence alignment of squid hnRNPA/B-like Proteins 2 with human hnRNPA1 protein showed the sequences in the N-terminal half were highly conserved in the hnRNPA/B subtype, and sequences in the C-terminal half has a conserved *steric zipper motif* with an overlay in the M9 domains. The Clustal W program was used to sequences align (Altschul SF et al., 1990), and legend infograms is indicated left below b) The beta amyloid fibers shown the incorporation of Thioflavin-T (Th-T) fluorescent dye when was measured directly in plates with excitation wavelength of 412 nm. Each 200 μl of samples containing: rP2 (30 μg) in 4 M urea and 1 M urea; S1 (45 μg) in 1 M urea; rp2 / S1 in 1 M urea in phosphate buffer, pH 7.5, at r.t. with 15 seconds of stirring before each dosing time fluorescence spectra of incorporated into amyloid fibrils. The beta amyloid fibers formation in the Aβ peptide (80 μg in 200 μl) were used as positive control of the incorporation of Th-T fluorescent c) The distinct phosphorylation between dimer (p65) and monomer (p37) forms. SDS-PAGE 10% gel stained with Coomassie Colloidal Blue (CCB) and Phosphoprotein Gel Stain (PGS, Invitrigen), shows tissue-purified p65 and p37 by Ionic exchange and reverse phase chromatography (p65 RP and p37 RP), the bacterially expressed and purified recombinant hnRNPA/B-like protein 2 by Ni2+-affinity chromatography (rP2). The Casein (α/β Casein) and Albumin bovine (BSA) proteins were used as positive control. The positions of the immunoreactive bands corresponding to p65, p37 are indicated and asterix is any other protein

### Squid hnRNPA/B-like Protein 2 relocation to stress granules

We have previously shown that squid hnRNPA/B-like Proteins 2 share a high degree of structural homology with human hnRNPA1 and hnRNPA2/B1 (Lopes et al., 2019). This similarity suggests cellular properties conserved between them. To investigate subcellular distribution of hnRNPA/B-like Protein 2 and human hnRNPA1 protein in a neuronal context, we used neuroblastoma cells (SH-SY5Y). SH-SY5Y cells were transfected with pEGFP-C1 vector containing the ORF of hnRNPA/B-like 2 expressing rP2 fused to GFP. The recombinant rP2-GFP expression is limited to the nucleous, showing a granular and interchomatinic location, being excluded from nucleoli. A similar labeling pattern is shown for anti-hnRNPA1 antibody. Regions of colocalization are seen in the nucleous and also shows some nuclear regions of more intense staining (Fig 4).

**Figure 4.**
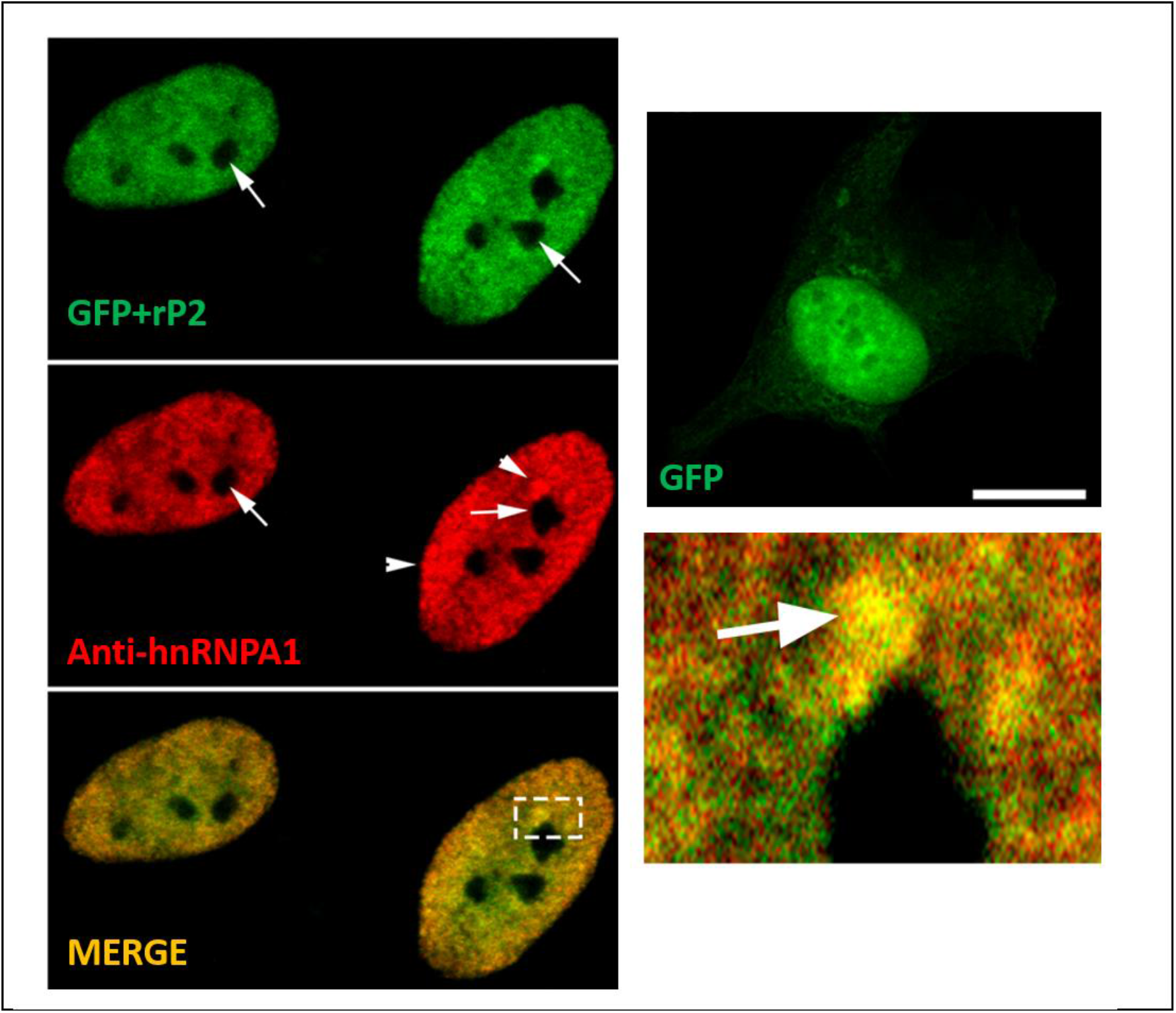
rP2 fused to GFP is exclusively nuclear. Human SH-SY5Y cells were transfected with pEGFP-C1 vector containing the ORF of hnRNPA/B-like 2 expressing rP2 fused to GFP (GFP+rP2) or pEGFP-C1 vector alone (GFP, left insert). The recombinant expression of GFP+rP2 is limited to the nucleous, showing a granular labelling excluded from nucleoli (arrows on the left figures). A similar labeling pattern is shown for anti-hnRNPA1 antibody (red). Regions of colocalization are seen in the nucleous and also shows some nuclear regions of more intense staining (see arrowheads). A nuclear colocalization region was amplified in right insert (arrows). Scale bar: 10 μm

SH-SY5Y cells were also transfected with pmCherry-C1 vector containing the ORF that codifies squid hnRNPA/B-like Protein 2 (mCherry-rP2) and double-labeled with monoclonal anti-Poly(A) binding protein (PABP) and anti-hnRNPA1 antibodies. SH-SY5Y cells expressing mCherry-rP2 was osmotically stressed by 400 mM sorbitol, which lead to the formation of cytoplasmic stress granules (SGs) containing PABP, rP2 and human hnRNPA1 (Fig 5a). After one hour of recovery in a medium without sorbitol, the amount of SGs containing rP2 (mCherry-rP2) and hnRNPA1 protein decreased in cytoplasm (Fig 5a; 1h recovery). PABP protein exhibited slower movement and less complete recovery than hnRNP A1 and rP2 (Fig 5b).

**Figure 5.**
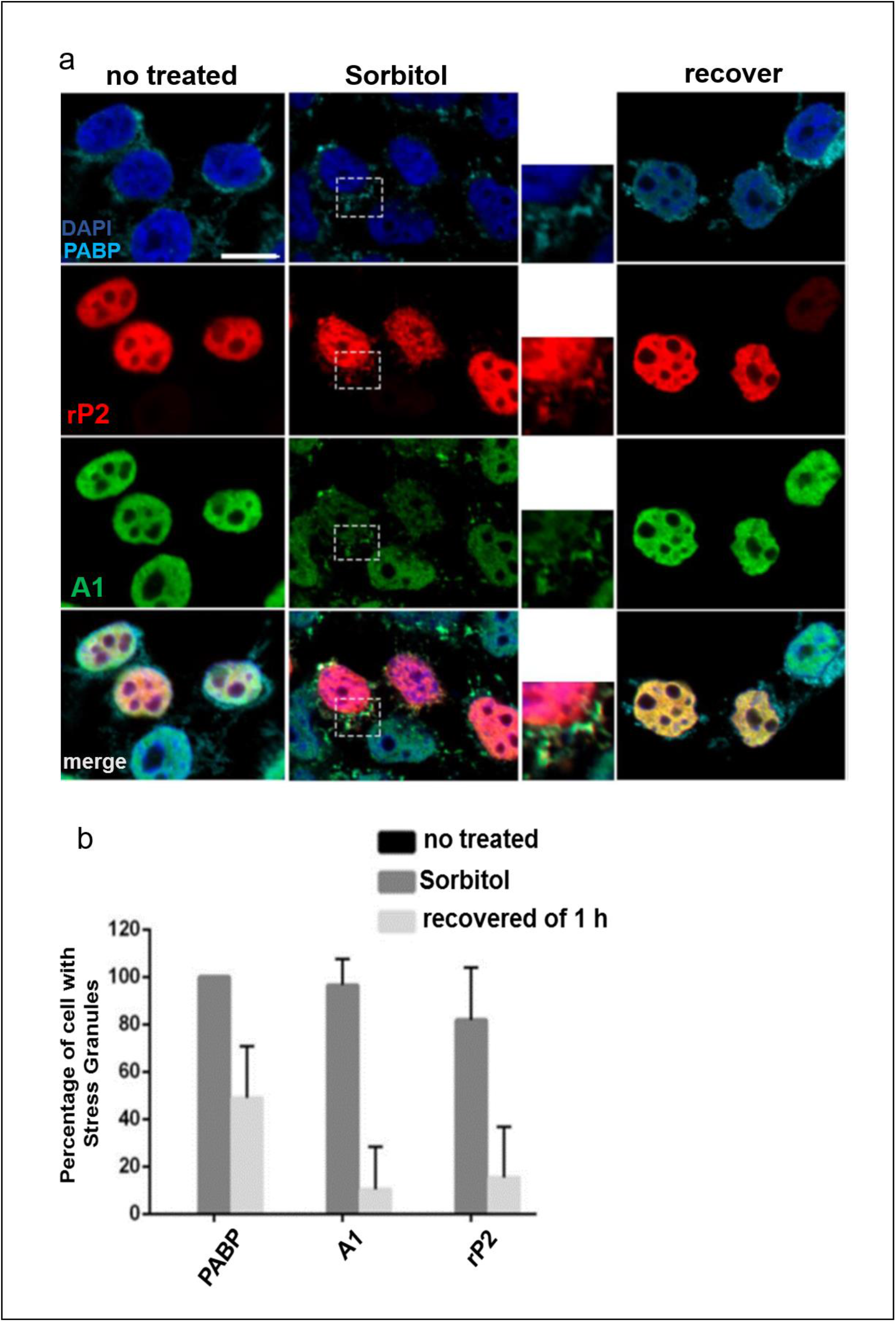
Sorbitol induced stress granules containing PABP, rP2 and human hnRNPA1. a) SH-SY5Y cells were transfected with mCherry-C1 vector containing the ORF of squid hnRNPA/B-like Protein 2 (red) and labeled with anti-PABP (cyan) and anti-human hnRNPA1 (green). The nuclei were stained with DAPI (blue). Hyperosmolar stress was induced by 400 mM sorbitol treatment for 1h, which led the formation of cytoplasmic stress granules (dashed line boxes) containing PABP, rP2 and hnRNPA1. After 1 h treatment with sorbitol, cells were washed in culture medium to remove sorbitol and allowed to recover, for 1h b) Percentage of cells expressing mCherry-rP2 in cytoplasmic granules including hnRNPA1 and PABP in each condition were calculated and plotted in a bar graph. About forty cells were counted for each condition analyzed. Analyses were performed on GraphPad Prism software. Experiments were performed using a Zeiss inverted microscope LSM 780 AxioObserver. Scale bar 10 μm

### Squid hnRNPA/B-like Protein 2 dynamicity in osmotic stress-activated cells

Stress granules (SGs) are dynamic structures that undergoes fusion, fission, and flowing in the cytosol. When cell reestablishes its normal physiological condition, SGs are disassembled and the proteins retained in the granules, as well as messenger RNAs, return to their site of action. In this way, messenger RNAs may be available again for translation (Protter & Parker, 2016). GFP-rP2 dynamics in SH-SY5Y cells under osmotic stress was observed by Fluorescence recovery after photobleaching (FRAP) assay. Data suggest that in sorbitol-stressed cells, GFP-rP2 moves in and out of SGs very quickly, with a fluorescence recovery rate of approximately 80% and a recovery time (measured in Ʈ / 2) of approximately 16 seconds (Fig. 6a and b). In non-stressed cells the recovery rate of nuclear GFP-rP2 is approximately 90%, and the recovery time is somewhat less, Ʈ / 2 = 10 seconds (Fig. 6 a and b). The data are consistent with process that driven dynamic of interactions with high-density RNP complexes in stress granules (SGs) at the cytoplasm (Lindquist, 1981, Kedersha et al., 1999; Guil S et al., 2006, Kedersha et al., 2013; Molliex et al., 2015; Banani S.F et al., 2016; Wheeler et al., 2016; Protter & Parker, 2016).

**Figure 6.**
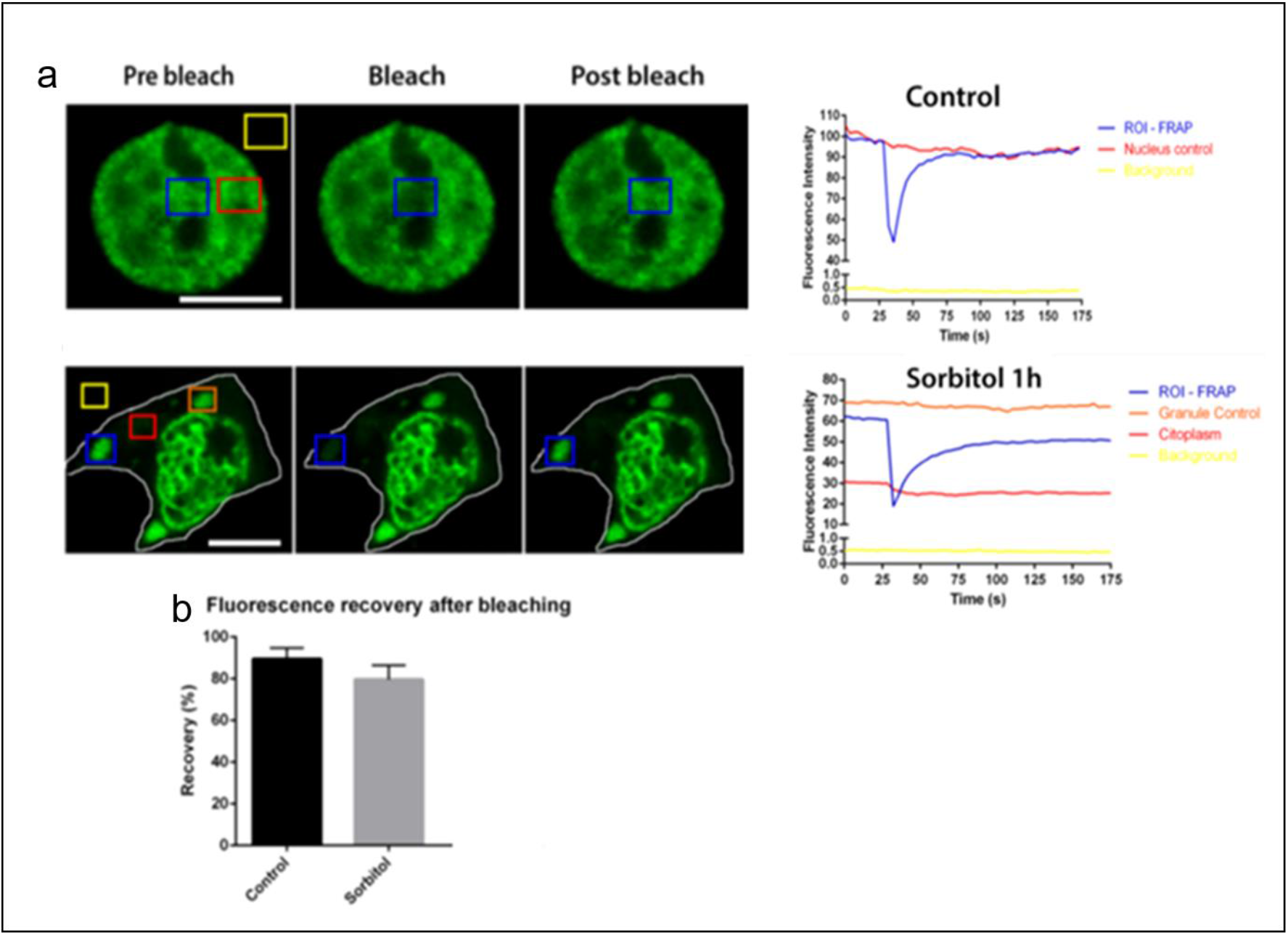
GFP-rP2 dynamicity in SHSY-5Y cells under osmotic stress observed by FRAP. Cells expressing GFP-rP2 were subjected to FRAP under control conditions, under cellular stress caused by treatment with 400 mM Sorbitol, for 1h. In each experiment, the regions of interest where FRAP was performed are delineated by the blue square (ROI-FRAP). As internal controls, red, orange and yellow squares, had their fluorescence intensity measured without bleaching a) Typical cells are shown, and the time lapse graph of fluorescence intensity of a FRAP assay where ROI-FRAP confined a nuclear region. The red square covered an adjacent nuclear region. The yellow square was located out of the nucleus, where no fluorescence is observed (background). On cells treated with sorbitol, ROI-FRAP covered a putative cytoplasmic stress granule, while the orange square marked an independent cytoplasmic granule, the red square, a cytoplasmic region without granules, and the yellow square the intercellular background. The cytoplasm is delimited by a white line b) Percentage of fluorescence recovery after bleaching was calculated as described in Methods. The values from six cells for each condition analyzed were averaged and plotted here + SEM. Analyses were performed on Orange Data Mining software, and the graphics on GraphPad Prism software. FRAP were performed on a Zeiss inverted microscope LSM 780 AxioObserver. Scale bars 10 μm

## Discussion

The monomeric (p37) and dimeric (p65) forms of squid hnRNPA/B-like Protein 2 are found in squid optic lobes associated with RNA molecules and co-sedimented with hnRNPA3, hnRNPA1, hnRNPA1-like 2 and ELAV-like in a heterogeneous nuclear ribonucleoprotein complex (Lopes et al.,2020). The large number of highly conserved of RNA-binding proteins allow many different combinations of hnRNPs are involved in packaging of nascent mRNAs into granules in the cells. An understanding of biochemical mechanisms involved in the formation and stability of the dimeric form of hnRNPA/B-like Protein 2 may bring insights in the dynamic protein–protein and RNA–protein binding into RNA granules.

In previously studies, was proposed a structural model of squid recombinant hnRNPA/B-like Protein 2(rP2) that supported the formation of a stable homodimer that, in part, was maintained from hydrogen and electrostatic bonds. The N-terminal of rP2 has two RNA binding domains or RRM domains linked by a loop, such that the beta sheets are oriented toward the surface to form the sites of interaction with RNA molecules (Lopes et al., 2019). These sites of interaction with RNA molecules are formed by disulfide bonds between RNA binding domains in a poly(A)-binding protein can regulate the ability of this protein to bind mRNA (Fong C. L et al., 2000).

In this scenario, we evidenced that squid recombinant hnRNPA/B-like Protein 2(rP2) can be interacting with squid RNA molecules, when inhibited the precipitation of rP2 in low urea solution (Fig. 2 a, cuvet 6 and Fig. S1). However, when optic lobe extract (S1) was pre-treated with ribonuclease A (RNase A) followed from incubation with the rP2, it leads to the formation of higher SDS-stable dimers (≥ 80-kDa) (Fig. 2c, cuvet 5). Remarkably, an amount increase of SDS-stable dimers formation can be visualized with high DTT (Fig. 2d). These data suggest that disulfide bonds formation between RRM domains of rP2 and squid RNAs could be regulated the availability of monomers of rP2 involved in the formation of the dimeric form, however endogenous factor(s) in the extract produce post translational alterations in dimers of rP2, that induce its dimerization giving the SDS-stable forms *in vitro*.

Notably, the sequence alignment of squid hnRNPA/B-like protein 2 and human hnRNP A1 protein primary structure; showed differences from Serine (14/42) and Cysteine (4/2); aromatic (26/20) and charged (18/14) aminoacids residues numbers, respectively. Our attention was caught to localization of four Cysteines aminoacids residues in the N-terminal half of squid hnRNPA/B-like protein 2 and a differences number of phospho-aminoacids potential residues (Fig. 3a).

So, we showed the mobility of recombinant hnRNPA/B-like Protein 2 (rP2) by gel filtration chromatography; where was noticed a disrupting their oligomers in two distinct fractions (~80-kDa and ~40-kDa), when the sample was treated with high concentrations of DTT, suggesting a involvement of disulfide bonds formation between RRM domains of rP2 visualized by Urea–PAGE (Fig. 1a and 1b). Note that to solubilize and purify hnRNPA/B-like protein 2, the buffers required high concentrations of urea (Lico et al., 2015), it could dramatically change the N-terminal of squid hnRNPA/B-like protein 2 (globular region), lead the formation of disulfide bounds between rP2 to transform into a Urea-stable dimers. However, to transform into SDS-stable dimers in low urea, has be appropriate post-translational modifications Its not clear exactly, as endogenous factor(s) in the extract induce its dimerization, although one attractive idea is that, post-translational modifications could change the structure of the hnRNPA/B-like protein 2 to revealed their C-terminal (disordered region), in which contain critics domains (steric zipper motif) that could be involved its dimerization, giving an SDS-stable conformation. For it, the hnRNPA/B-like Protein 2 showed structures conserved in critics domains such a steric zipper motif (Fig. 3 a and b), having a high degree of structural homology to the hnRNPA/B subfamily of RNA binding (Kim HJ et al., 2013). In addition, we observed a distinct phosphorylation between dimer (p65) and monomer (p37), its come from their cellular behavior into squid optic lobe (Fig. 3c). Taking over the high degree of structural homology that squid hnRNPA/B-like Protein 2 shares with human hnRNPA1 protein (Lopes et al., 2019), suggest a cellular function conserved between these proteins.

Here, we investigated the subcellular distribution of hnRNPA/B-like Protein 2 and human hnRNPA1 protein in a neuronal context, where both accumulates predominantly in the nucleus and excludes nucleolar regions in human SH-SY5Y cells, see Fig 4. On the other hand, when cultured SH-SY5Y cells undergo hyperosmotic stress, under treatment with sorbitol, rP2 displays a punctate pattern in the cytoplasm colocalizing with hnRNPA1 and PABP into stress granules (SGs), see Fig 5. Following the dynamic of SGs undergo fusion, fission and can shuttle out of the nucleus and locate to cytoplasmic (Fig. 6)., as Guil S. et al., 2006, suggesting a conserved cellular property of squid and human hnRNPA/B family proteins. So, the propensity to aggregate reversibly is an important property for components of RNA granules (Banani S.F et al., 2016; Protter & Parker, 2016). However, this important property may have the dire consequence of facilitating uncontrolled aggregation, particularly when genetic risk factors are present (Polymenidou M. and Cleveland D. W., 2017; Ramaswami M et al., 2013).

In summary, our data indicate a conserved structure and cellular properties of squid and human hnRNPA1 proteins and shed light on biochemical mechanisms of the dimeric form of rP2. Remarkably, hnRNPA/B-like protein 2 showed two molecular species of dimers; from mobility in urea–PAGE, suggesting a involvement of disulfide bonds formation between RRM domains (globular region) of rP2, in which may be involves the binds from RNA molecules, and SDS-resistant dimers visualized on SDS–PAGE; which suggests that endogenous factor(s) could induces posttranslational modifications in hnRNPA/B-like protein 2 that promote the formation of the SDS-stable dimers, its could involves the C-terminal (disordered region), in which contain critics domains (steric zipper motif). An understanding of the transition between monomeric and dimeric forms molecular bring insights on inclusion body formation in neuropathologies.

## Supporting information

https://drive.google.com/file/d/1J6cl43NjYHx_L9Wr-QhrPY1EI16pzcL5/view?usp=sharing

## Abbreviations

rP2: recombinant hnRNPA/B-like protein 2
hnRNP: heterogeneous nuclear ribonucleoprotein
ORF: open reading frame
PAGE: polyacrylamide gel electrophoresis
PBS: phosphate buffered saline
RNP: ribonucleoprotein
RNP1/ RNP2: core sequences of RNA recognition motifs
SDS: sodium dodecyl sulfate
SG: stress granules
BFS: bovine fetal serum
FRAP: Fluorescence Recovery After Photobleach

## Acknowledgements and contributions

DTPL (Performed experiments biochemical and Prepared manuscript); GSL (Performed experiments immunofluorescence); MLPL (Analyzed data) and REL (designed the experiments); received financial support from the Fundação de Amparo à Pesquisa do Estado de São Paulo (FAPESP), the Conselho Nacional de Desenvolvimento Científico e Tecnológico (CNPq) and the Fundação de Apoio ao Ensino, Pesquisa e Assistência do Hospital das Clínicas da FMRP-USP (FAEPA). Special thanks to Silvia Andrade and Domingos Pitta for expert technical help, and M.Sc Elizabete R. Milani for technical help with confocal microscopy, performed at Laboratório Multiusuário de Microscopia Confocal—LMMC, Fapesp 2004/08868-0

## References

Altschul SF, Gish W, Miller W, Myers EW, Lipman DJ (1990) Basic local alignment search tool. J Mol Biol, 215:403–410.

Chomczynski, P.; Sacchi, N. Single–step method of RNA acid guanidinium thiocyanate-phenol-chloroform extraction. Anal Biochem, 162:156–159 1987.

Lico DTP, Rosa JC, DeGiorgis JA, deVasconcelos EJR, Casaletti L, Tauhata SBF, Baqui MMA, Fukuda M, Moreira JE, Larson RE (2010) A novel 65 kDa RNA-binding protein in squid presynaptic terminals. Neurosci, 166:73–83.

Lico DT, Lopes GS, Brusco J, Rosa JC, Gould RM, De Giorgis JA, Larson RE (2015) A novel SDS-stable dimer of a heterogeneous nuclear ribonucleoprotein at presynaptic terminals of squid neurons. Neuroscience, 300, 381–392.

Lopes, G. S.; Lico, D.T.P; Rocha, R. S.; Oliveira, R. R.; Sebollela, A. S.; Paco-Larson, M. L.; Larson, R. E. (2019) A phylogenetically conserved hnRNP type A/B protein from squid brain. Neuroscience Letters, 696: 219–224

Maris C, Dominguez C, Allain FH (2005) The RNA recognition motif, a plastic RNA-binding platform to regulate post-transcriptional gene expression. FEBS J, 272:2118–2131.

W.M. Michael, M. Choi, G. Dreyfuss (1995) A nuclear export signal in hnRNP A1: a signal mediated, temperature-dependent nuclear protein export pathway. Cell, 83 (3) 415–422.

Siomi H., G. Dreyfuss (1995) A nuclear localization domain in the hnRNP A1 protein. J.Cell Biol, 129 (3) 551–560.

Guil, J.C. Long, J.F. Cáceres (2006) hnRNPA1 relocalization to the stress granules reflects a role in the stress response. Mol. Cell. Biol, 26 (15) 5744–5758.

Maharana S, Wang J, Papadopoulos DK, Richter D, Pozniakovsky A, Poser I, Bickle M, Rizk S, Guillén-Boixet J, Franzmann TM, Jahnel M, Marrone L, Chang YT, Sterneckert J, Tomancak P, Hyman AA, Alberti S (2018) RNA buffers the phase separation behavior of prion-like RNA binding proteins. Science, 360:918–921.

He, Y. and Smith, R. (2009) Nuclear functions of heterogeneous nuclear ribonucleoproteins A/B. Cellular and Molecular Life Sciences, 1239–1256

Dreyfuss G, Kim VN, Kataoka N (2002) Messenger-RNA-binding proteins and the messages they carry. Nat Rev Mol Cell Biol, 3:195–205

Lopes S. Gabriel., Brusco Janaina, Rosa C José., Larson E. Roy & Lico T. P Diego. (2020) Selectively RNA interaction by a hnRNPA/B-like protein at presynaptic terminal of squid neuron. Invertebrate Neuroscience, 20:1–14

Yalda Baradaran-Heraviab, ChristineVan Broeckhovenab and Julievan der Zee (2020) Stress granule mediated protein aggregation and underlying gene defects in the FTD-ALS spectrum Neurobiology of Disease. 134:104639

Douglas N Joshua., Lidia A. Gardner, Hannah E Salapa, and Michael C. Levin (2016) Antibodies to the RNA Binding Protein Heterogeneous Nuclear Ribonucleoprotein A1 Colocalize to Stress Granules Resulting in Altered RNA and Protein Levels in a Model of Neurodegeneration in Multiple Sclerosis. J Clin Cell Immunol, 7(2): 402

Elvira George, Sylwia Wasiak, Vanessa Blandford, Xin-Kang Tong, Alexandre Serrano, Xiaotang Fan, Maria del Rayo Sánchez-Carbente, Florence Servant, Alexander W. Bell, Daniel Boismenu, Jean-Claude Lacaille, Peter S. McPherson, Luc DesGroseillers and Wayne S. Sossin (2006) Characterization of an RNA Granule from Developing. BrainMolecular & Cellular Proteomics, (4) 635–651

Purice MD and Taylor JP (2018) Linking hnRNP Function to ALS and FTD Pathology. Front. Neurosci, 12:326.

McDonald, K.K., Aulas, A., Destroismaisons, L., Pickles, S., Beleac, E., Camu, W., Rouleau, G.A. Vande Velde, C (2011) TAR DNA-binding protein 43 (TDP-43) regulates stress granule dynamics via differential regulation of G3BP and TIA-1. Hum. Mol. Genet, 20, 1400–1410.

Langdon, E.M., Qiu, Y., Ghanbari Niaki, A., McLaughlin, G.A., Weidmann, C.A., Gerbich, T.M., Smith, J.A., Crutchley, J.M., Termini, C.M., Weeks, K.M., Myong, S., Gladfelter, A.S (2018) mRNA structure determines specificity of a polyQ-driven phase separation. Science, 360: 922–927.

Van Treeck, B., Protter, D.S.W., Matheny, T., Khong, A., Link, C.D., Parker, R. (2018) RNA self-assembly contributes to stress granule formation and defining the stress granule transcriptome. Proc. Natl. Acad. Sci. U. S. A, 115: 2734–2739.

Zhang, K., Daigle, J.G., Cunningham, K.M., Coyne, A.N., Ruan, K., Grima, J.C., Bowen, K.E., Wadhwa, H., Yang, P., Rigo, F., Taylor, J.P., Gitler, A.D., Rothstein, J.D., Lloyd, T.E (2018) Stress granule assembly disrupts nucleocytoplasmic transport. Cell, 173:958–971 e917.

Polymenidou Magdalini, Don W Cleveland (2017) Biological Spectrum of Amyotrophic Lateral Sclerosis Prions. Cold Spring Harb Perspect Med, 7:a024133

Ramaswami, M., Taylor, J.P., Parker, R. (2013) Altered ribostasis: RNA-protein granules in degenerative disorders. Cell, 154:727–736.

Kim HJ, Kim NC, Wang Y-D, Scarborough EA, Moore J, et al. (2013) Mutations in prion-like domains in hnRNPA2B1 and hnRNPA1cause multisystem proteinopathy and ALS. Nature, 495:467–473.

McLellan T (1982) Electrophoresis buffers for polyacrylamide gels at various pH. Anal Biochem, 126:94–99.

Molliex, A., Temirov, J., Lee, J., Coughlin, M., Kanagaraj, A.P., Kim, H.J., Mittag, T., Taylor, J.P (2015) Phase separation by low complexity domains promotes stress granule assembly and drives pathological fibrillization. Cell, 163: 123–133.

Protter, D.S.W., Rao, B.S., Van Treeck, B., Lin, Y., Mizoue, L., Rosen, M.K., Parker, R (2018) Intrinsically disordered regions can contribute promiscuous interactions to RNP granule assembly. Cell Rep, 22:1401–1412.

Protter David S. W. and Roy Parker (2016) Principles and Properties of Stress granules. Trends Cell Biol, 26(9): 668–679.

Lindquist, S., 1981. Regulation of protein synthesis during heat shock. Nature, 293:311–314.

Kedersha, N.L., Gupta, M., Li, W., Miller, I., Anderson, P (1999) RNA-binding proteins TIA-1 and TIAR link the phosphorylation of eIF-2 alpha to the assembly of mammalian stress granules. J. Cell Biol, 147:1431–1442

Kedersha, N., Ivanov, P., Anderson, P (2013) Stress granules and cell signaling: more than just a passing phase? Trends Biochem. Sci, 38:494–506.

Wheeler, J.R., Matheny, T., Jain, S., Abrisch, R., Parker, R (2016) Distinct stages in stress granule assembly and disassembly. Elife5 Trends Biochem. Sci, 38: 494–506.

Fong L Christopher., Amy Lentz, and Stephen P. Mayfield (2000) Disulfide Bond Formation between RNA Binding Domains Is Used to Regulate mRNA Binding Activity of the Chloroplast Poly(A)-binding Protein. The Journal of Biological Chemistry, 275:8275–8278.

Kwiatkowski Jr., T.J., Bosco, D.A., Leclerc, A.L., Tamrazian, E., Vanderburg, C.R., Russ, C., Davis, A., Gilchrist, J., Kasarskis, E.J., Munsat, T., Valdmanis, P., Rouleau, G.A., Hosler, B.A., Cortelli, P., de Jong, P.J., Yoshinaga, Y., Haines, J.L., Pericak-Vance, M.A., Yan, J., Ticozzi, N., Siddique, T., McKenna-Yasek, D., Sapp, P.C., Horvitz, H.R., Landers, J.E., Brown Jr., R.H. (2009) Mutations in the FUS/TLS gene on chromosome 16 cause familial amyotrophic lateral sclerosis. Science, 323:1205–1208

Liu, Y.C., Chiang, P.M., Tsai, K.J (2013) Disease animal models of TDP-43 proteinopathy and their pre-clinical applications. Int. J. Mol. Sci, 14:20079–20111.

Liu, Q., Shu, S., Wang, R.R., Liu, F., Cui, B., Guo, X.N., Lu, C.X., Li, X.G., Liu, M.S., Peng, B., Cui, L.Y., Zhang, X (2016) Whole-exome sequencing identifies a missense mutation in hnRNPA1 in a family with flail arm ALS. Neurology, 87:1763–1769.

Mackenzie, I.R., Nicholson, A.M., Sarkar, M., Messing, J., Purice, M.D., Pottier, C., Annu, K., Baker, M., Perkerson, R.B., Kurti, A., Matchett, B.J., Mittag, T., Temirov, J., Hsiung, G.R., Krieger, C., Murray, M.E., Kato, M., Fryer, J.D., Petrucelli, L., Zinman, L., Weintraub, S., Mesulam, M., Keith, J., Zivkovic, S.A., Hirsch-Reinshagen, V., Roos, R.P., Zuchner, S., Graff-Radford, N.R., Petersen, R.C., Caselli, R.J., Wszolek, Z.K., Finger, E., Lippa, C., Lacomis, D., Stewart, H., Dickson, D.W., Kim, H.J., Rogaeva, E., Bigio, E., Boylan, K.B., Taylor, J.P., Rademakers, R (2017) TIA1 mutations in amyotrophic lateral sclerosis and frontotemporal dementia promote phase separation and Alter stress granule dynamics. Neuron, 95:808–816 e809.

Aguzzi Adriano and Lawrence Rajendran (2009) The transcellular spread of cytosolic amyloids, prions, and prionoids. Neuron, 64(6):783–90.

Buratti E (2018) TDP-43 post-translational modifications in health and disease. Expert Opin Ther Targets, 22:279–293.

Banani S.F., Allyson M. Rice, William B. Peeples, Yuan Lin, Saumya Jain, Roy Parker, and Michael K. Rosen (2016) Compositional Control of Phase-Separated Cellular Bodies. Cell, 166:651–663.

Khong A., Tyler Matheny, Saumya Jain, Sarah F Mitchell, Joshua R Wheeler, and Roy Parker (2017) The stress granule transcriptome reveals principles of mRNA accumulation in stress granules. Mol Cell, 68:808–820.e5.

