## Supplementary material for "Biochemical and subcellular characterization of a squid hnRNPA/B-like protein in osmotic stress activated cells reflects molecular properties conserved in this protein family": https://drive.google.com/file/d/1J6cl43NjYHx_L9Wr-QhrPY1EI16pzcL5/view?usp=sharing

**Supplement information**

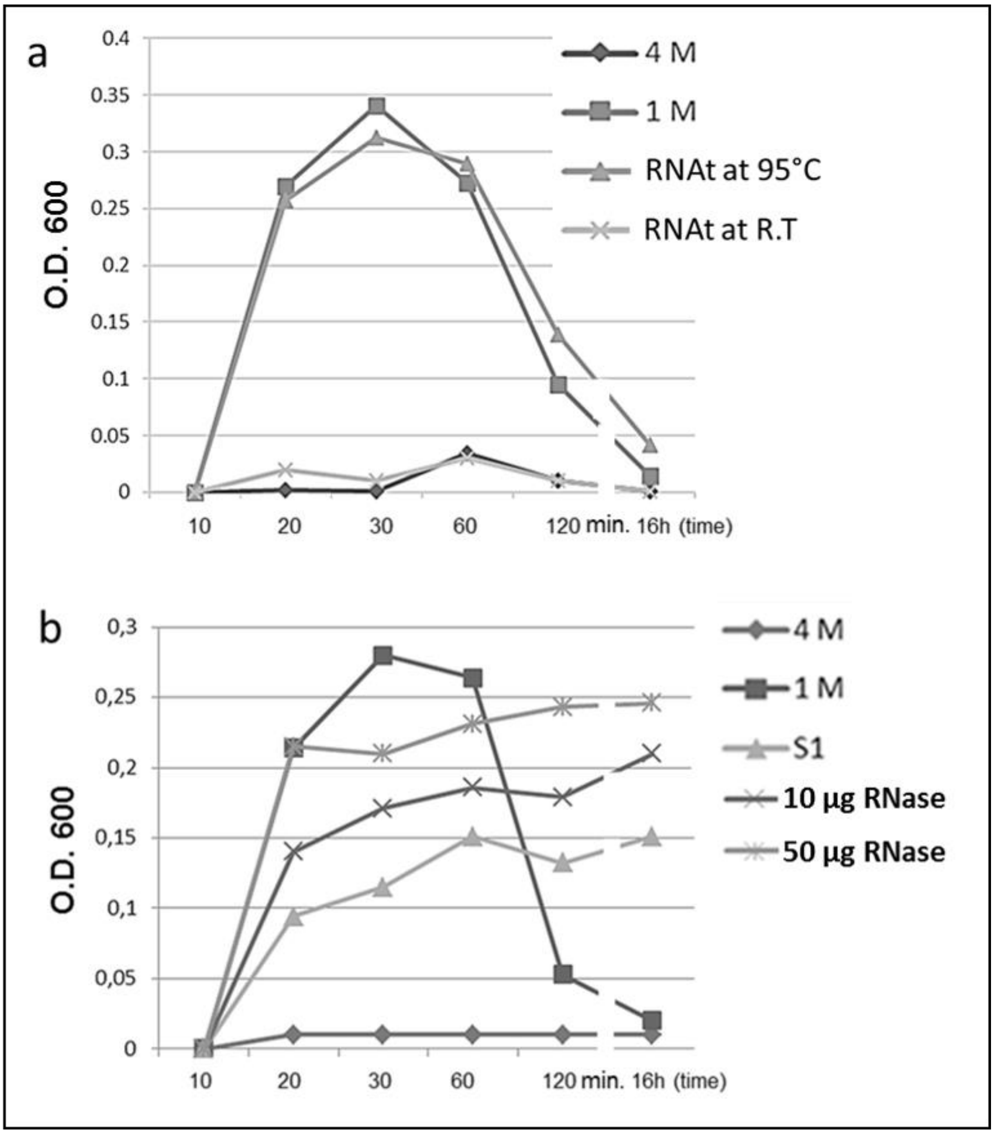


Fig S1

**Fig. S1 - The precipitation of rP2 show chromatograms in 1M urea monitored by light scattering at 600 nm for 16 hours at room temperature (r.t)**

a) Reactions were carried out directly in plastic cuvets in 50 mM Phosphate Buffer, pH 7.5, 100 mM NaCl, 25 mM DTT, 30 mM Imidazole, 0,5 mM ATP and 1 mM protease inhibitors (aprotinin; Benzamidine and pefabloc). All cuvets contain 200 µg/ml of rP2 in Phosphate Buffer. Line ( ) rP2 in 4 M urea; line ( ) rP2 in 1 M; line ( ) rP2/RNA at 95 °C in 1 M and line ( ) rP2/RNA at r.t in 1 M. Squid total RNA isolated from optic lobe add 40 U / ml out RNA. To denaturation secondary structure one fraction with 25 µg/ml RNA was pre-incubated at 95 °C for 5 minutes and other sample fraction at r.t, to incubated with rP2 *in vitro* experiments

b) Precipitation of rP2 in 1M urea for 16 hours at room temperature (r.t). Reactions were carried out directly in plastic cuvets in 50 mM Phosphate Buffer, pH 7.5. All cuvets contain 200 µg/ml of rP2 in 50 mM Phosphate Buffer. Line ( ) rp2 in 4 M urea; line ( ) rp2 in 1 M;line ( ) rp2/S1 in 1 M. Supernatants 1 (S1) from squid optic lobe were treatments by 30 minutes with line ( )10 and line ( ) 50 µg/mlRNase

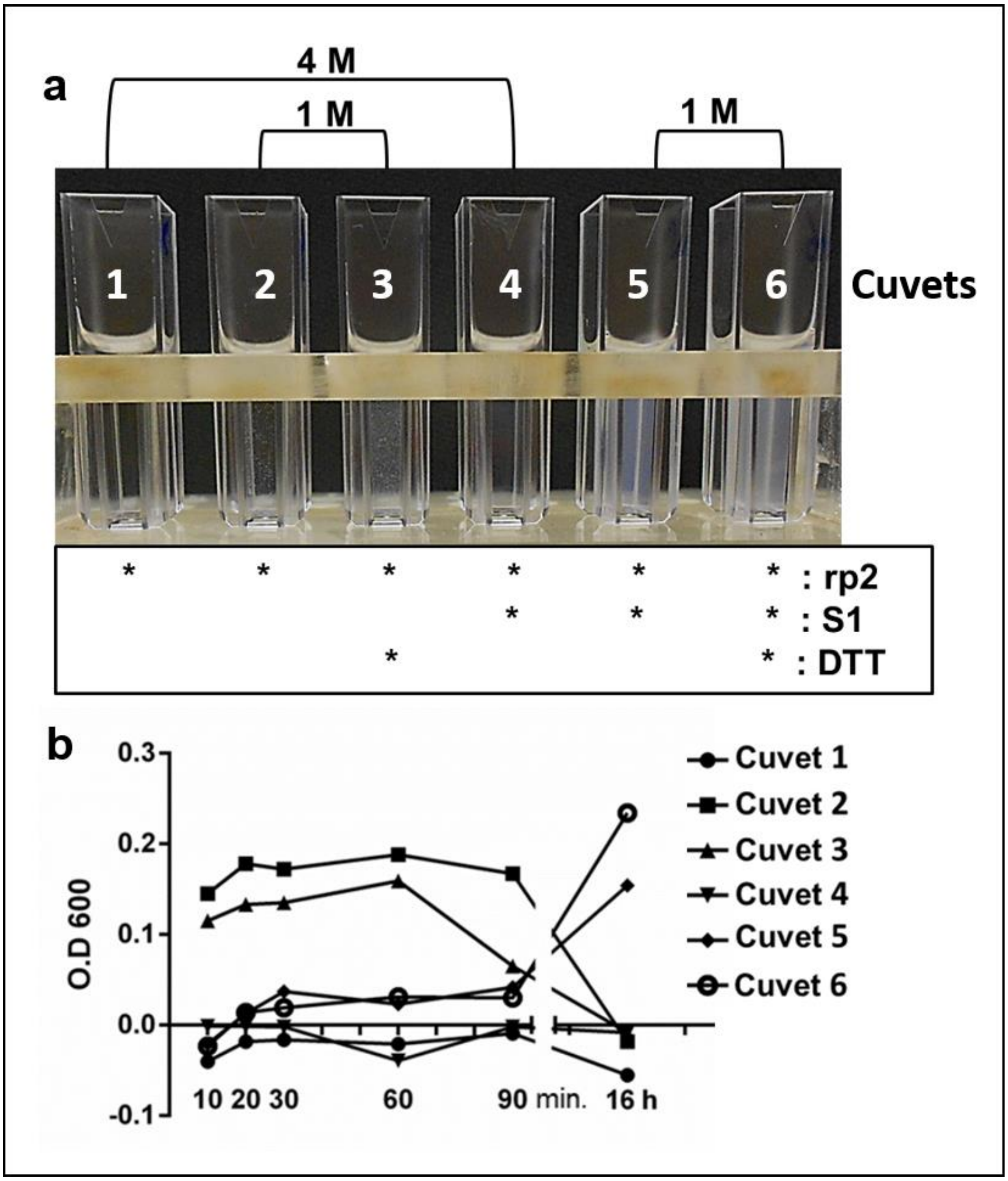
a
4 M
1 M
1 M
1
2
3
4
5
6
Cuvets
* * * * * * : rp2
* * * : S1
* * : DTT
b
0.3
0.2
0.1
0.0
-0.1
O.D 600
10 20 30
60
90 min.
16 h
Cuvet 1
Cuvet 2
Cuvet 3
Cuvet 4
Cuvet 5
Cuvet 6

Fig S2

**Fig. S2 - The precipitation of recombinant hnRNP A / B - like protein 2(rp2) in the presence of 4 M and 1 M urea**

a) The sample was carried out directly in plastic cuvets in 50 mM Phosphate Buffer, pH 7.5, 100 mM NaCl, 25 mM DTT, 30 mM Imidazole, 0,5 mM ATP and 1 mM protease inhibitors (aprotinin; Benzamidine and pefabloc). All cuvets contain 200 µg/ml of rP2 in 50 mM Phosphate Buffer. Cuvet 1 and 4 with 4 M urea; cuvet 2,3,5 and 6 with 1 M urea. Asterix indicated the addition of 220 µg/ml of S1 from squid optical lobe in and plus 125 mM of DTT in Phosphate Buffer

b) The precipitation of rP2 show chromatogram in all condition described in (a) and monitored by light scattering at 600 nm for 16 hours at room temperature (r.t)

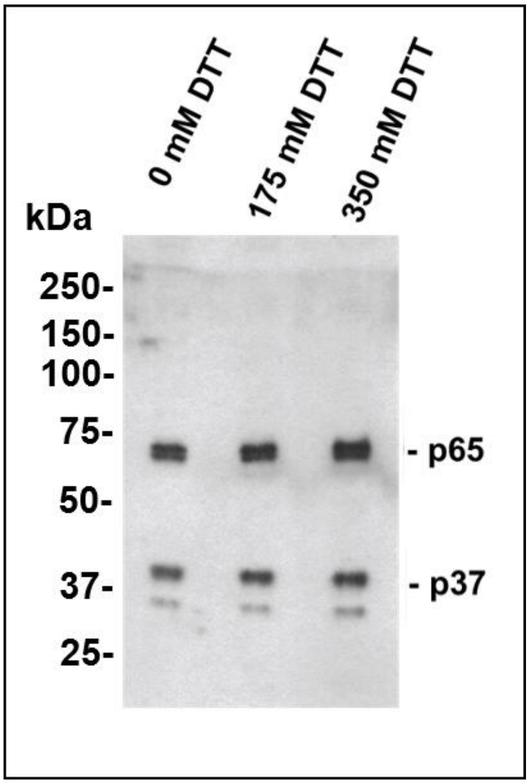


Fig S3

**Fig.S3 – The endogenous p65 is resistant the high concentrations of DTT**. Western blots from squid optic lobes extract was extracted in sample buffer at different concentrations of DTT and denatured for 5 min. at 95°C. Immunoblot revealed with α-sqRNP2 polyclonal antibody. The positions of the immunoreactive bands corresponding to p65, p37 are indicated
